# It takes two (Las1 HEPN Endoribonuclease Motifs) to cut the RNA right

**DOI:** 10.1101/760058

**Authors:** Monica C. Pillon, Kevin H. Goslen, Jason G. Williams, Robin E. Stanley

## Abstract

Las1 is an essential endoribonuclease that is well-conserved across eukaryotes and a newly established member of the HEPN (higher eukaryotes and prokaryotes nucleotide-binding) nuclease family. HEPN nucleases participate in diverse RNA cleavage pathways and share a short HEPN nuclease motif important for RNA cleavage. While most HEPN nucleases participate in stress activated RNA cleavage pathways, Las1 plays a fundamental role in processing the pre-ribosomal RNA. Underscoring the significance of Las1 function, mutations to the *LAS1L* gene have been associated with neurological dysfunction. Two juxtaposed Las1 HEPN nuclease motifs create its composite nuclease active site, however the roles of the individual HEPN residues are poorly defined. Here we show through a combination of *in vivo* and *in vitro* studies that both Las1 HEPN nuclease motifs are required for nuclease activity and fidelity. Through in-depth sequence analysis and systematic mutagenesis, we define the consensus Las1 HEPN nuclease motif and uncover its canonical and specialized elements. Using reconstituted Las1 HEPN-HEPN’chimeras, we define the molecular requirements for RNA cleavage. Intriguingly, both copies of the Las1 HEPN motif are necessary for nuclease specificity revealing that both HEPN motifs participate in coordinating the RNA within the active site. Taken together, our work reveals critical information about HEPN nuclease function and establishes that HEPN nucleases can be re-wired to cleave alternative RNA substrates.

## Text

Endoribonucleases play critical roles in the processing, maturation, and destruction of diverse RNAs by catalyzing the hydrolysis of phosphodiester bonds (1). The HEPN (higher eukaryotes and prokaryotes nucleotide-binding) superfamily is an emerging group of endoribonucleases spanning across all walks of life. The HEPN domain is characterized by a small α-helical domain originally suggested to be important for nucleotide binding (2). Subsequent bioinformatic analysis transformed the field through the widespread identification of HEPN-containing proteins and the classification of a catalytic-subset that encode for a short, but defined RϕxxxH (where ϕ is often N, D, or H, and x is any amino acid that can vary from 3 to 5 residues) endoribonuclease motif (3) (Fig. 1A). The expanding list of HEPN nucleases highlights their profound role in numerous biological processes, most notably host-defense and stress response pathways. For example, the eukaryotic HEPN nuclease Ire1 triggers the unfolded protein response and the bacterial HEPN nuclease SO_3166 is the toxic component of the dueling type II toxin-antitoxin system (4, 5). In contrast to these defense systems, the HEPN nuclease Las1 ensures the translational capacity of the cell through its role in ribosome assembly (6). Furthermore, the bacterial Cas13 CRISPR effectors represent a set of programmable HEPN-containing nucleases that are gaining broad attention for their potential therapeutic applications (7–11). Despite the strong prevalence of the HEPN domain in critical RNA-targeting nucleases, the molecular basis for how HEPN nucleases catalyze RNA cleavage has remained elusive.

**Fig. 1.**
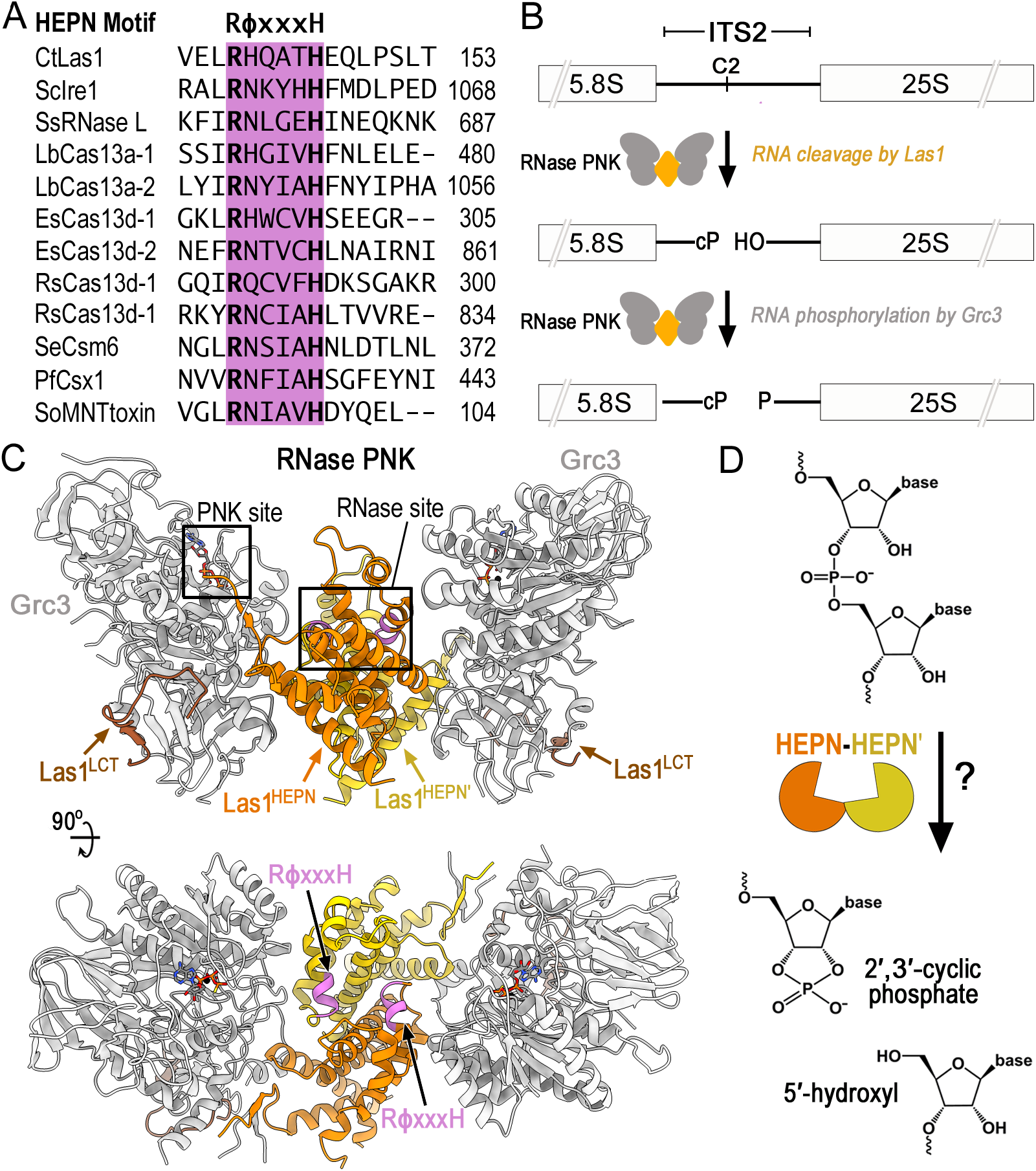
Members of the HEPN superfamily encode a canonical RϕxxxH motif responsible for RNA cleavage. (*A*) Amino acid sequence alignment of the HEPN motif from several different HEPN nucleases. The HEPN canonical motif RϕxxxH (where ϕ is commonly N, H or D and x is any residue) is responsible for RNA cleavage and highlighted in purple. Abbreviations are as follows: *Chaetomium thermophilum* (Ct), *Saccharomyces cerevisiae* (Sc), *Sus Scrofa* (Ss), *Leptotrichia buccalis* (Lb), *Eubacterium siraeum* (Es), *Ruminococcus species* (Rs), *Staphylococcus epidermidis* (Se), *Pyrococcus furiosus* (Pf) and *Shewanella oneidensis* (So). (*B*) Diagram of ITS2 pre-rRNA processing by RNase PNK. RNase PNK is composed of the Las1 RNase and the Grc3 PNK. Las1 cleaves the C2 site leaving a 2’,3’-cyclic phosphate (cP) and a 5’-hydroxyl (OH) (14). Subsequently, Grc3 phosphorylates (P) the 5’-hydroxyl end marking the ITS2 for decay by a series of exoribonucleases. (*C*) Orthogonal views of *C. thermophilum* RNase PNK model (PDB ID 6OF3). Grc3 promoters are show in gray and Las1 protomers are shown in orange and yellow. Las1 RϕxxxH motifs are highlighted in purple and ATP-γS are shown as sticks. Las1 RNase and Grc3 PNK active sites are boxed. (*D*) Cartoon schematic of metal-independent RNA cleavage by members of the HEPN superfamily. HEPN members dimerize (orange/yellow) to form an active nuclease, which cleaves the phosphodiester backbone through an unclear mechanism resulting in a 2’,3’-cyclic phosphate and 5’-hydroxyl RNA ends.

The Las1 HEPN nuclease harbors unique characteristics that distinguishes it from other members of the HEPN superfamily. Defects in Las1 (Las1L in humans) are linked to motor neuron diseases and X-linked intellectual disability (12, 13), however its molecular function in ribosome production was only elucidated following its recent classification as an HEPN nuclease (3, 14). Unlike most HEPN nucleases that are activated following cellular stress (3), Las1 is essential for cell growth and proliferation (6, 15). Las1 plays a fundamental role in processing precursor ribosomal RNA (pre-rRNA) (6, 14–17). Ribosome production involves a complex and highly coordinated pre-rRNA processing cascade tasked with removing four spacer sequences from the pre-rRNA (14, 18–21). Removal of the ITS2 (internal transcribed spacer 2), which lies between the 5.8S and 25S rRNAs, is initiated by Las1 pre-rRNA cleavage at the defined C2 site (Fig. 1B) (14). Following endoribonucleolytic cleavage, the ITS2 is phosphorylated by the Grc3 RNA kinase, which in turn triggers the recruitment of 5’- and 3’-exoribonucleases that sequentially degrade the ITS2 spacer (14, 16, 22). Further distinguishing Las1 from other HEPN nucleases is Las1’s dependence on its binding partner, the Grc3 polynucleotide kinase for nuclease activation (23). While no other HEPN nuclease is known to require an auxiliary protein for HEPN activation, Las1 and Grc3 are reliant on one another for protein stability and enzyme activation (15, 23, 24). Together the ribonuclease (RNase) Las1 and polynucleotide kinase (PNK) Grc3 assemble into a tetrameric complex, known as RNase PNK. Through this higher-order assembly, RNase PNK coordinates its dual enzymatic functions through a poorly understood mechanism of molecular cross-talk that ensures efficient processing of the pre-rRNA (23–25).

Las1 adopts a canonical HEPN active site suggesting it shares a common mechanism for RNA cleavage as other HEPN nucleases. We recently solved a series of cryo-electron microscopy structures of RNase PNK, which unveiled its butterfly-like architecture (26). Two Las1 protomers homodimerize at the axis of symmetry where the HEPN domains form the ‘body’ of the butterfly, while a Grc3 protomer makes up each ‘wing’ and holds the HEPN domains together (Fig. 1C) (26). This is reminiscent to all characterized HEPN nucleases, which require dimerization of two HEPN domains to activate nuclease activity (4, 5, 7, 27, 28). Furthermore, the composite Las1 HEPN active site resembles that of other HEPN nucleases, which are formed by the juxtaposition of two RϕxxxH HEPN motifs (Fig. 1C) As with all members of this superfamily, the Las1 HEPN nuclease catalyzes metal-independent RNA cleavage resulting in the production of a terminal 2’,3’-cyclic phosphate and 5’-hydroxyl (Fig. 1D). Despite the extensive structural and biochemical characterization of numerous HEPN nucleases, the role of the individual residues comprising HEPN RϕxxxH motifs remains unclear for this superfamily. With the widespread prevalence and functional significance of HEPN nucleases in diverse biological processes, including the use of HEPN-associated CRISPR-Cas nucleases for *in vivo* RNA-targeting applications (7, 11, 29–31), it is paramount to define the molecular mechanism of HEPN domains in RNA cleavage.

To gain a better understanding of the molecular features driving RNase PNK nuclease activity, we generated a series of Las1 RϕxxxH HEPN variants to determine the functional significance of each individual residue. Through a combination of *in vivo* studies in *Saccharomyces cerevisiae* and *in vitro* activity assays, we determined that the flanking invariant arginine and histidine residues along with several intervening residues of the RϕxxxH motif are critical for nuclease activity. Unexpectedly, we also discovered that the Las1 RϕxxxH HEPN motif contributes to nuclease fidelity and alteration of this motif leads to the generation of off-target RNA cleavage products. Thus, this work lays the foundation for engineering HEPN nucleases with altered specificity.

## Results and Discussion

### The Las1 HEPN Domain Encodes a Conserved RHxhTH Motif

Las1 is conserved across eukaryotes and encodes a six amino acid RϕxxxH motif within its HEPN domain. Previous work has established that the invariant arginine and histidine residues of RϕxxxH are critical to support *S. cerevisiae* (Sc) Las1 RNA cleavage *in vitro* and *in vivo* (14, 23, 24), yet little is known about the intervening residues comprising this motif. To determine the significance of the entire Las1 RϕxxxH HEPN motif, we curated over 300 unique Las1 orthologs from species spanning fungi to vertebrates and carried out a comprehensive sequence alignment. Analysis of the alignment encompassing the Las1 HEPN domain revealed a single well-conserved RϕxxxH motif previously associated with Las1 nuclease activity (Fig. 2A). This led to the discovery that beyond the first arginine (R1) and last histidine (H6) residues found in all HEPN family nucleases, the intervening residues within the Las1 RϕxxxH HEPN motif are also well-conserved. The second position within this motif is an invariant histidine (H2), which fits with the general trend that HEPN nucleases often encode for a polar amino acid at this position (3). On the other hand, the third position is a poorly conserved residue (x3) with a subtle preference for alanine and the fourth position has a strong preference for a hydrophobic amino acid (h4). While a subset of plant Las1 homologs encode for a serine residue at the fifth position, the majority of Las1 homologs harbor a threonine (T5). Based on this high sequence conservation, we define the consensus Las1 HEPN motif as RHxhTH (where x is any residue and h is a hydrophobic residue) (Fig. 2A).

**Fig. 2.**
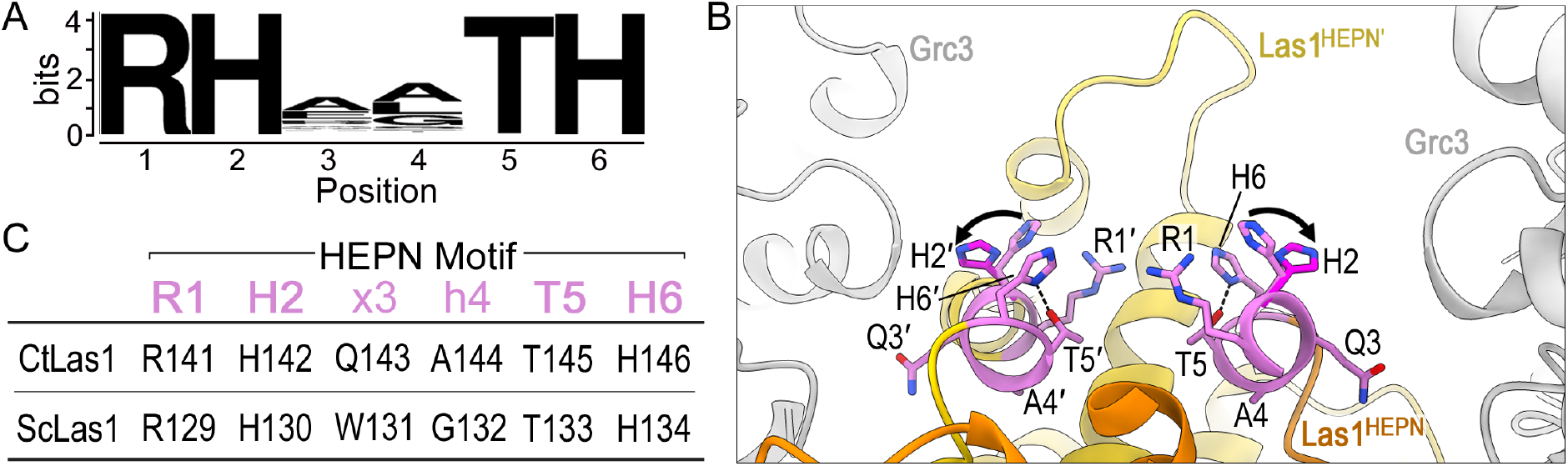
The Las1 HEPN nuclease encodes a consensus RHxhTH catalytic motif. (*A*) Sequence logo for the Las1 HEPN motif generated from over 101 vertebrates and 219 fungi orthologs. The height of each letter is correlated with its conservation within the motif. The logo defines the consensus Las1 HEPN motif RHxhTH (where x is any residue and h is commonly hydrophobic). (*B*) Las1 HEPN nuclease active site from *C. thermophilum* RNase PNK in state 1 (PDB ID 6OF3) and state 2 (PDB ID 6OF2). Grc3 protomers are colored in gray and the two Las1 HEPN domains are shown in orange and yellow. The second HEPN protomer is denoted by prime. Residues of the Las1 HEPN motifs RHxhTH are shown in purple. Conserved residues R1, H2, T5 and H6 all face the catalytic center whereas x3 (CtLas1 Q3) points towards the cleft formed between Las1 and Grc3 and h4 (CtLas1 A4) contributes to the hydrophobic core. Black arrows highlight the rearrangement of invariant residue H2 between state 1 (light purple) and state 2 (dark purple). Dotted line between residues T5 and H6 represents a putative hydrogen bond. Residues 108-140 of the proximal Las1^HEPN^ protomer (orange) was removed for display purposes. (*C*) Table of the equivalent RHxhTH consensus residues in *C. thermophilum* Las1 (CtLas1) and *S. cerevisiae* Las1 (ScLas1).

The structural architecture surrounding the Las1 RHxhTH motif reveals the molecular basis for its residue preference at each position. Our recent cryo-electron microscopy reconstructions of *Chaetomium thermophilum* (Ct) RNase PNK revealed the juxtaposed Las1 RHxhTH motifs within the composite nuclease active site (Fig. 2B) (26). Each Las1 RHxhTH motif is encoded within an α-helix that lies at the interface of the Las1 HEPN homodimer. By capturing multiple conformational states of RNase PNK, we revealed that H2 is an important active site switch that toggles between nuclease active and inactive conformations (26). In the active state, H2 is pointed towards the center of the active site whereas H2 is pointing away in the inactive state (Fig. 2B). While H2 undergoes a distinct conformational rearrangement, the remaining residues of the Las1 RHxhTH motif remain largely unchanged. In the nuclease active conformation, the four well-conserved residues of the RHxhTH motif (R1, H2, T5, H6) all point towards the nuclease active site suggesting they each play an important role in catalysis (Fig. 2B-C). The well-conserved T5 lies at the base of the nuclease active site where it is within hydrogen bonding distance of H6 and could contribute to the spatial positioning of H6. The other two residues, which lie within the middle of the motif, are positioned away from the active site. The variable residue at the third position (CtLas1 Q3) points towards the broad cleft formed between the nuclease and kinase active sites of RNase PNK explaining the lack of conservation at this position. Conversely, the hydrophobic residue at the fourth position (CtLas1 A4) is buried in the Las1 HEPN-HEPN’core and explains its strong preference for a non-polar residue to preserve its native fold (Fig. 2B-C). Together, the identification of the Las1 HEPN consensus RHxhTH motif, along with the recent structural characterization of RNase PNK, uncovers the molecular requirements for the Las1 composite HEPN nuclease active site.

### The Las1 RHxhTH Motif is Essential for *S. cerevisiae* Cell Proliferation

To investigate the functional significance of the Las1 RHxhTH motif, we disrupted the motif and monitored its functional effects in *S. cerevisiae* (Sc) (15, 23). Analogous to all Las1 homologs, ScLas1 is composed of a well-conserved N-terminal HEPN nuclease domain followed by a poorly conserved coiled-coil (CC) domain and a conserved C-terminal tail, herein called LCT for Las1 C-terminal Tail (Fig. 1C and 3A) (6, 17). The ScLas1 HEPN domain harbors the HEPN nuclease motif ^129^RHWGTH, which emulates the consensus RHxhTH motif found in all Las1 homologs. While previous work has established that ScLas1 variants harboring R1A, H2A, or H6A mutations are non-viable in *S. cerevisiae* and unable to cleave the ITS2 pre-rRNA *in vitro* (14, 26);it is unknown if any other amino acid substitutions are tolerated within this motif. To determine the requirements of the RHxhTH consensus motif, we utilized a strain of *S. cerevisiae* containing a tetracycline-inducible promoter (tetO_7_) upstream of endogenous *LAS1 (tet-LAS1)* (26). This strain was modified to include a 3x-Myc tag on the N-terminus of endogenous *GRC3 (tet-Las1/Myc-GRC3)* so that we could monitor Grc3 expression. Addition of doxycycline (DOX) to the growth medium represses the expression of endogenous Las1 and inhibits cell growth and proliferation (Fig. 3B) (15). The *tet-LAS1* and *tet-LAS1/Myc-GRC3* strains were transformed with a plasmid harboring a 3x-Flag tagged wild-type Las1 construct (Flag-Las1) or an empty ARS1-CEN4 YCplac vector (32). We tested the complementation of Flag-Las1, by repressing endogenous *LAS1* expression with doxycycline at 30**°**C. Unlike the transformed yeast strain harboring the empty vector, yeast expressing Flag-Las1 rescues *S. cerevisiae* growth in the presence of doxycycline both on solid media and in liquid culture (Fig. 3B-C).

**Fig. 3.**
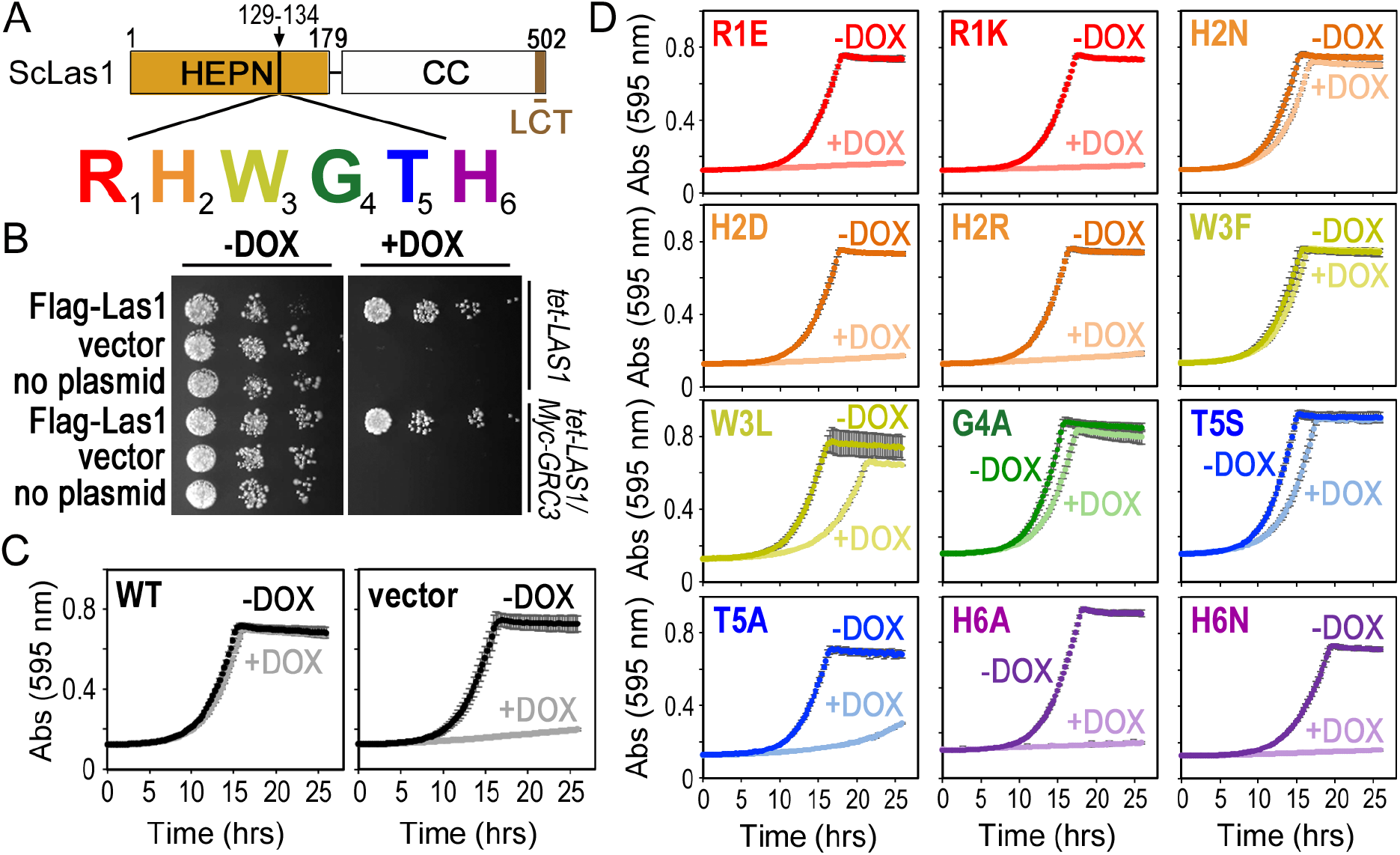
The Las1 RHxhTH motif is essential for *S. cerevisiae* proliferation. (*A*) Cartoon of *S. cerevisiae* (Sc) Las1 with the numbers defining the amino acid residue domain boundaries. The Las1 HEPN domain is shown as an orange box, the Coiled-coil (CC) domain is a white box and the Las1 C-terminal tail (LCT: residues 469-502) is shown in brown. Individual residues of the ScLas1 RHxhTH motif (^129^RHWGTH) are highlighted in red, orange, yellow, green, blue and purple, respectively. (*B*) *S. cerevisiae tet-LAS1* and *tet-LAS1/3xMyc-GRC3* strains were either left untransformed (no plasmid) or transformed with plasmids encoding wild type Las1 with an N-terminal 3x Flag tag (Flag-Las1) and empty YCplac 33 vector (vector). Serial dilutions of each strain were grown on YPD agar in the absence (-DOX) and presence (+DOX) of 40 μg/ml doxycycline at 30°C for 3 days. Doxycycline represses transcription of endogenous *LAS1*. (*C*) Growth curves of *tet-LAS1/3xMyc-GRC3* strains transformed with control plasmids encoding wild type Flag-Las1 (WT) and empty YCplac 33 (vector). Strains were incubated at 30°C in the absence and presence of doxycycline and absorbance was recorded over 25 hours at 595 nm. Each curve is an average of three independent replicates and error bars define the standard deviation. (*D*) Growth curves as seen in panel *C* with the *tet-LAS1/3xMyc-GRC3* strain transformed with Flag-Las1 RHxhTH variants (see Table S2). Color scheme is the same as seen in panel *A*.

To assess the requirements of the Las1 RHxhTH motif, we engineered a series of single missense mutations to each residue (Fig. 3A and Table S1). Twelve individual Las1 RHxhTH HEPN motif variants (R1E, R1K, H2N, H2D, H2R, W3F, W3L, G4A, T5S, T5A, H6A, and H6N) were transformed into the *tet-LAS1/Myc-GRC3* strain (Table S2). The complementation of these variants was tested by repressing endogenous *LAS1* expression with doxycycline at 30°C and growth curves were recorded over a 25 hour time period by measuring the absorbance at 595 nm (Fig. 3D). Disruption of the first arginine (R1E, R1K) and last histidine (H6A, H6N) residues prevents cell proliferation underscoring the significance of these residues for Las1 function. Mutations to the invariant H2 residue (H2N, H2D, H2R) yielded variable results. The H2D and H2R variants are non-viable, however the H2N variant only causes a minor growth defect. While the role of H2 in Las1 function is unknown, the observation that either a histidine or asparagine can support Las1 function *in vivo* is suggestive of a role in RNA engagement and/or positioning in the nuclease active site. Disruption of either the third or fourth residue from the Las1 RHxhTH HEPN motif (W3F, W3L, G4A) only causes mild growth defects, which is not unexpected given the poor sequence conservation and position within the RNase PNK structure. Mutagenesis of the threonine at the fifth position to a serine (T5S) causes a mild growth defect, while an alanine (T5A) causes a severe growth defect. This corresponds well with a subset of plant Las1 homologs that encode a serine at this position and suggests these homologs are functional. A serine residue at the fifth position may be tolerated because it could still maintain a hydrogen bond to the adjacent catalytic histidine (H6), as seen with T5 in the RNase PNK structure (Fig. 2B). Conversely, an alanine substitution (T5A) would prevent hydrogen bonding to H6, thus potentially disrupting Las1 function and explaining its severe growth defect. Interestingly, the catalytic histidine found in the nuclease active site of RNase A also forms a similar hydrogen bond with a nearby aspartic acid residue. While disruption of this hydrogen bond leads to a modest defect in RNA cleavage, this interaction is thought to promote proper orientation of the catalytic histidine for RNA hydrolysis and enhance the conformational stability of the active site (33, 34). Therefore, these results suggest a threonine or serine at the fifth position is important for Las1 function. We also tested temperature sensitivity by plating serial dilutions of these strains on solid media and monitored their growth at five different temperatures in the presence and absence of doxycycline (Fig. S1). With the exception of W3L, T5S and T5A, the majority of the HEPN variants do not display temperature sensitivity. Temperatures higher or lower than 30°C result in more significant growth defects for the W3L, T5S and T5A variants. Taken together these results highlight the importance of the entire Las1 RHxhTH HEPN motif for cell viability in *S. cerevisiae*.

### Las1 RHxhTH Variants do not Disrupt Protein Stability or Grc3 Association

We performed quality controls to determine that the growth defects observed with the Las1 HEPN variants are not the result of Las1 instability. To confirm that the HEPN variants do not compromise Las1 protein stability *in vivo*, we grew *tet-LAS1/Myc-GRC3* strains expressing the Las1 HEPN variants to mid-log phase in the presence of doxycycline at 30**°**C and then analyzed the whole cell lysate by Western blot. Due to the co-dependence of Las1 and Grc3 for protein stability (15), we also analyzed endogenous levels of 3xMyc-tagged Grc3 (15). Flag-tagged Las1 was detected with an anti-Flag antibody, Grc3 was detected with an anti-Myc antibody and tubulin was used as a loading control. Expression of the Las1 HEPN variants does not substantially alter the endogenous levels of Las1 or Grc3 *in vivo* (Fig. S2A). Therefore, the growth defects caused by the Las1 HEPN variants are not due to a loss in Las1 protein stability.

The Las1 HEPN variants also retain their association with Grc3 and support higher-order assembly of the RNase PNK complex. The Las1 HEPN nuclease directly depends on its binding partner Grc3 to execute its ITS2 pre-rRNA processing activity (23). To ensure that the defects observed in cell growth and proliferation are not the result of an inability to associate with Grc3, we monitored assembly of RNase PNK using the Las1 HEPN variants and wild type Grc3. To reconstitute recombinant RNase PNK complexes, we generated a series of *Escherichia coli* co-expression vectors encoding wild-type Grc3 along with poly-histidine tagged Las1 HEPN variants (Table S3). Using affinity chromatography, we immobilized Las1 HEPN variants and assessed their association with Grc3. SDS-PAGE analysis of the purified samples revealed that all of the Las1 HEPN variants stably expressed and co-purified with Grc3 (Fig. S2B). Since the higher-order assembly of the RNase PNK complex is also crucial for ITS2 pre-rRNA processing (23), we monitored RNase PNK hetero-tetrameric assembly by gel filtration. All of the RNase PNK variants show similar retention volumes to wild-type RNase PNK suggesting they all maintain their hetero-tetrameric organization (Fig. S2C). Thus, this confirms that the Las1 HEPN variants do not hinder Grc3 association or oligomerization of RNase PNK.

### Las1 RHxhTH Motif is Required for C2 pre-rRNA Cleavage

The Las1 HEPN variants reveal the importance of the RHxhTH motif for C2 cleavage. We speculated that the yeast growth defects observed in the presence of the Las1 HEPN variants were due to a defect in C2 pre-rRNA cleavage. To determine the effects of the Las1 variants on C2 pre-rRNA cleavage, we performed *in vitro* C2 cleavage assays using a 3’-fluorescently labeled ITS2 RNA mimic (C2 RNA substrate) (Fig. 4A). These reactions were carried out under enzyme excess, where RNase PNK variants were in molar excess over the C2 RNA substrate. Incubation of the C2 RNA substrate with wild-type RNase PNK results in a specific 5’-hydroxyl RNA fragment that can be resolved on a denaturing urea gel and was previously mapped to the C2 site (Fig. 4B) (23). In contrast, incubation of the C2 RNA substrate with the RNase PNK Las1 variants led to the observation that many of the variants are deficient at C2 cleavage. A representative gel summarizing C2 cleavage reactions of all the HEPN variants is shown in Fig. 4B. The variants that cause severe growth defects in *S. cerevisiae* (R1E, R1K, H2D, H2R, T5A, H6A, H6N) are unable to cleave the C2 site indicating that the observed growth defects are the result of an inability to cut the pre-rRNA (Fig. 4B-C). In contrast, the Las1 HEPN variants that cause minor growth defects (H2N, W3L, G4A, T5S) are able to cleave the ITS2 at the C2 site, albeit less efficiently than wild-type (Fig. 3D and 4C).

**Fig. 4.**
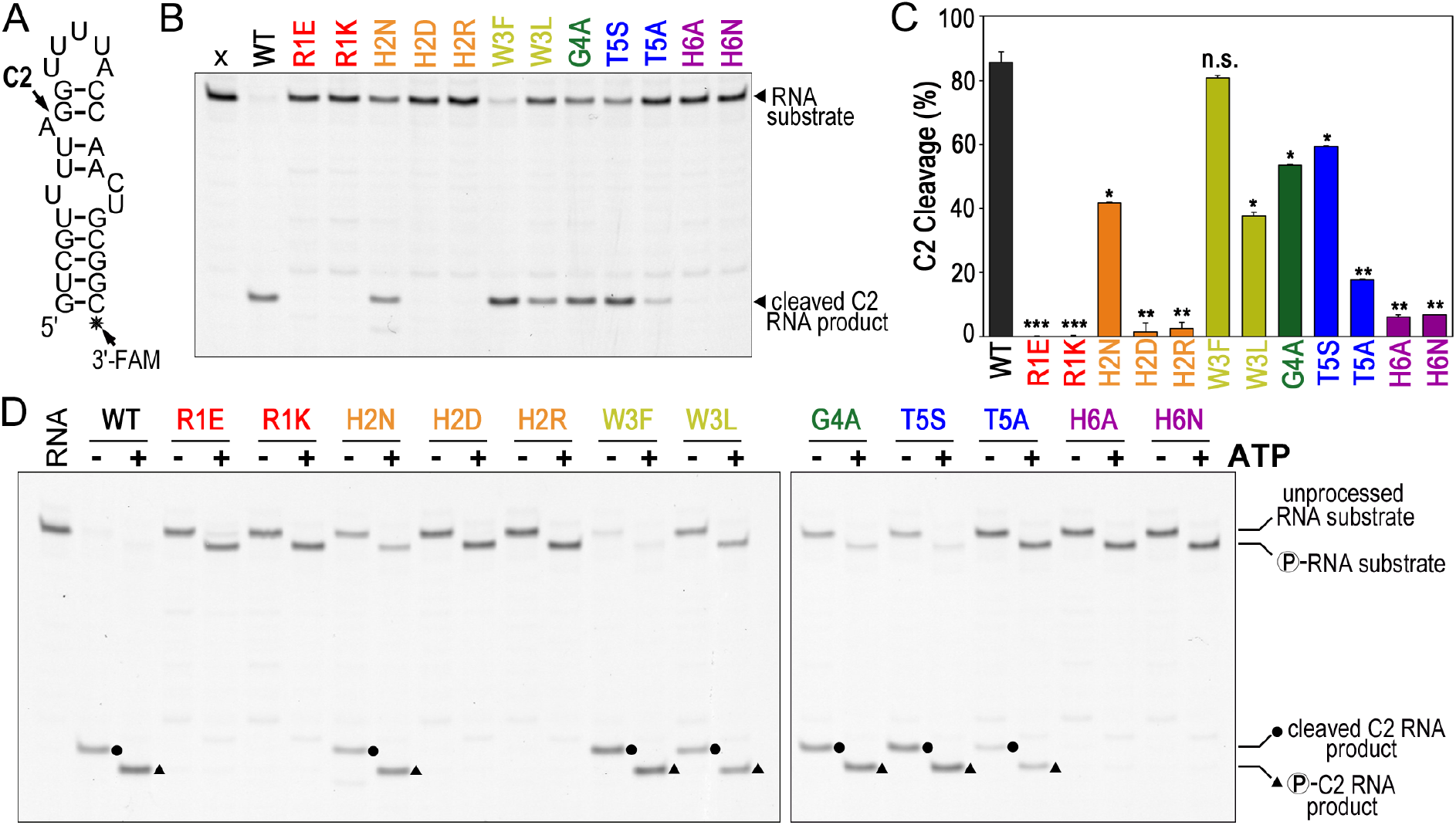
Las1 RHxhTH is critical for efficient C2 cleavage. (*A*) C2 RNA substrate used for *in vitro* C2 processing assays. This RNA substrate mimics the *S. cerevisiae* ITS2 C2 site (23). Black star marks the 3’-fluorescein label (3’-FAM). (*B*) Representative C2 RNA cleavage by recombinant *S. cerevisiae* Las1 RHxhTH variants bound to wild type Grc3. Excess ScRNase PNK variants (4.5 μM) were incubated with C2 RNA substrate (50 nM) for 1 hour at 37°C and resolved using denaturing urea gels. (*C*) Quantification of C2 RNA cleavage product shown in panel *B*. The mean and standard deviation were calculated from three independent replicates. n.s. not significant, \**P* < 6 x10^−3^, \*\**P* < 8 x10^−4^, and \*\*\**P* < 6 x10^−6^ were calculated by two-tailed Student’s *t* tests. (*D*) Representative C2 RNA cleavage and phosphorylation by recombinant ScRNase PNK variants. Protein variants (4.5 μM) were incubated with C2 RNA substrate (50 nM) in the absence (-) and presence (+) of 2 mM ATP for 1 hour at 37°C. RNA marks the reaction set in the absence of protein. Dots define C2 RNA cleavage products and triangles label C2 RNA phosphorylation products.

Beyond its role in nuclease activity, Las1 is also important for supporting nuclease-directed kinase activity, raising the possibility that the Las1 HEPN variants may also hinder Grc3 kinase activity. Previously we showed that the Las1^R1E,H6A^ variant hinders Grc3 phosphotransferase activity *in vitro* (23). To determine if the individual Las1 variants impact Grc3 kinase activity, we added ATP to the C2 cleavage reactions and visualized RNA phosphorylation through altered RNA mobility in a denaturing urea gel. All of the Las1 HEPN variants are able to phosphorylate either the 5’-hydroxyl end of the unprocessed C2 RNA substrate and/or the 5’-hydroxyl end of the C2 cleavage product (Fig. 4D). The observation of multiple phosphorylation events confirms our earlier work that showed the Grc3 kinase component of RNase PNK has broad RNA specificity (24). Taken together, these results indicate that the Las1 HEPN variants do not prevent Grc3 kinase activity on the ITS2 substrate, but are crucial for supporting Las1 nuclease activity.

### Both Las1 HEPN Motifs are Required for Nuclease Fidelity

Next, we sought to determine the significance of each individual Las1 RHxhTH motif for ITS2 pre-rRNA recognition and hydrolysis. Composite HEPN nuclease active sites are formed by two RϕxxxH motifs through either trans or cis assembly. Trans-homodimerization of HEPN domains is typically observed in HEPN nucleases, including RNase PNK, Ire1, RNase L, and Csm6 (5, 23, 27, 28, 35, 36). Cis-heterodimerization of tandem HEPN domains encoded within a single protomer has thus far only been observed in Cas13 nucleases (7, 11, 29, 31). Individual mutagenesis of the R1 or H6 residues from a single copy of the tandem HEPN heterodimer of the Cas13 subfamily impairs cleavage *in vitro* (11, 31). In contrast, trans complementation assays with HEPN homodimers of Ire1 and RNase L does not impair cleavage (28, 36). Collectively these results suggest that the requirement for juxtaposed R1 and H6 residues varies amongst HEPN family nucleases.

To determine whether RNase PNK requires both copies of its Las1 HEPN motifs for C2 cleavage, we engineered Las1 constructs that harbored mutations to a single copy of the HEPN homodimer. The constitutive dimerization of the Las1 HEPN domains poses a challenge for examining the significance of residues from the individual RHxhTH motifs. To overcome this problem, we used the RNase PNK structure to design a chimera of ScLas1 in which we fused two Las1 HEPN domains (HEPN|HEPN’) together with a flexible linker (Fig. 5A). The rational for the Las1 chimera was based off our previous work demonstrating that with the exception of the LCT, which is critical for Grc3 stability, the CC domain of Las1 is not required for RNase PNK nuclease and kinase activity *in vitro* (23). We reconstituted the chimeric ScRNase PNK complex using an *E. coli* co-expression vector encoding the Las1^HEPN|HEPN’^chimera, a polyhistidine tagged Las1^LCT^ and full-length Grc3 (Fig. 5A). First, we purified the chimeric RNase PNK complex and confirmed Las1^HEPN|HEPN’^retains its association with Grc3 and maintains its higher-order assembly using SDS-PAGE and gel filtration, respectively (Fig. S3A). Furthermore, we confirmed that the chimeric RNase PNK complex (WT|WT’) retains nuclease and kinase activities along with specificity for the C2 site (Fig. 5B: FL vs. WT|WT’and S3B). Chimeric RNase PNK titrations revealed that the Las1^WT|WT’^chimera cleaves the C2 RNA substrate almost as efficiently as full length Las1, while the double mutant Las1^H6A|H6A’^chimera is severely deficient in C2 cleavage (Fig. 5B and S3B). This confirms that the chimeric RNase PNK construct is a functional and C2-specific nuclease.

**Fig. 5.**
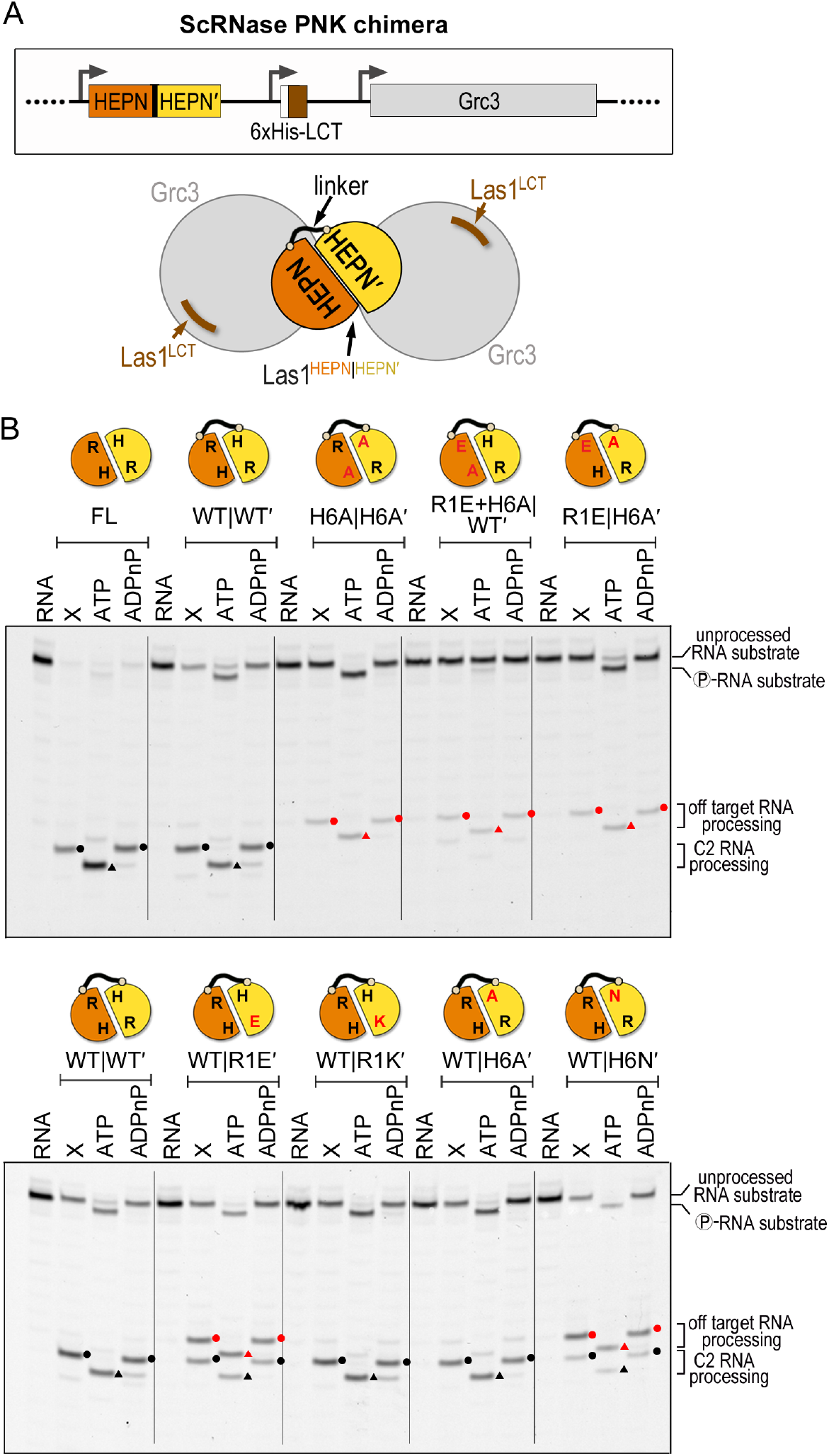
Both copies of the ScLas1 RHxhTH motif are required for C2 cleavage fidelity. (*A*) Cartoon schematic of the chimeric ScRNase PNK complex comprised of the Las1 HEPN|HEPN’dimer (orange|yellow) connected by a flexible linker, the Las1 LCT (brown) and wild type Grc3 (gray). The black line represents the linker tethering the Las1 HEPN|HEPN’dimer. The Las1 coiled-coil domain is dispensable for C2 RNA processing *in vitro (23)* and was removed to create the chimeric ScLas1^HEPN|HEPN’+LCT^-Grc3 complex. (*B*) Denaturing urea gels of RNA cleavage and phosphorylation reactions with full length RNase PNK (FL) and chimeric RNase PNK HEPN variants (HEPN|HEPN’). The lanes are marked as follows: RNA (100 nM RNA alone), X (0.8 μM RNase PNK variant and 100 nM RNA), ATP (0.8 μM RNase PNK variant, 100 nM RNA, and 10 mM ATP), ADPnP (0.8 μM RNase PNK variant, 100 nM RNA, and 10 mM ADPnP). Circles mark RNA cleavage products and triangles identify phosphorylated RNA. Black symbols specify C2 RNA processing products while red symbols are off-target RNA processing products.

After confirming that chimeric RNase PNK mimics the behavior of the full length complex *in vitro*, we generated a series of chimeric RNase PNK variants that encode missense mutations to a single RHxhTH motif. We carried out both nuclease and kinase assays under enzyme excess with double the amount of C2 RNA substrate to enhance the detection of low abundance RNA cleavage and phosphorylation products (Fig. 5B). Chimeric RNase PNK complexes harboring Las1^WT|R1E’^, Las1^WT|R1K’^, Las1^WT|H6A’^or Las1^WT|H6N’^all retained nuclease and kinase activity *in vitro*. This suggests that only three of the four R1, R1’, H6 and H6’residues are necessary for RNA hydrolysis and functional cross-talk with the Grc3 RNA kinase. Unexpectedly, several chimeric variants (WT|R1E’and WT|H6N’) also produced additional cleavage products (Fig. 5B, bottom gel). These additional products are all phosphorylated by Grc3 in the presence of ATP, but not a non-hydrolysable ATP analog, confirming that they must arise from Las1 cleavage. For instance, the chimeric RNase PNK comprising Las1^WT|R1E’^produces two cleavage products, one that coincides with canonical C2 cleavage and another that is the result of an off-target cleavage event which lies at least one nucleotide away from the C2 site. The chimeric RNase PNK complex harboring Las1^WT|H6N’^also produces the same two cleavage products, but the majority is the off-target product. To determine the identity of the off-target product, we performed liquid chromatography electrospray ionization mass spectrometry (LC-ESI-MS) analysis on the uncleaved C2 RNA substrate and the C2 RNA substrate following an incubation with chimeric RNase PNK harboring Las1^WT|H6N’^. The unprocessed C2 RNA substrate gave rise to a single peak above background that corresponds with the theoretical mass of the uncleaved C2 RNA substrate containing 5’- and 3’-hydroxyl ends (Fig. S4A). The mutant chimeric RNase PNK cleavage reaction gave rise to three RNA peaks above background including the unprocessed C2 RNA substrate and two product peaks (Fig. S4B). One product peak corresponds to an 8-nucleotide RNA fragment containing a 5’-hydroxyl and 2’,3’-cyclic phosphate while the second product is a 19-nucleotide RNA fragment containing 5’- and 3’-hydroxyl ends. This unambiguously maps the Las1-mediated off-target cleavage event to the phosphodiester bond 5’to the canonical C2 scissile phosphate (off-target site: U139-A140 of ScITS2). Thus, we confirm that the mutant chimeric RNase PNK complex has altered specificity and we denote the resulting off-target cleavage event as C2(−1) RNA cleavage. Taken together, this highlights the coordinated action of H6 and H6’, and to a less extent R1 and R1’, in maintaining C2 cleavage specificity.

To assess how strict the composite Las1 HEPN active site is for C2 specific RNA cleavage, we generated chimeric RNase PNK complexes harboring two mutations to the RHxhTH motif either in cis or in trans. As we previously observed with the Las1 double mutant H6A|H6A’, in cis (R1E+H6A|WT’) and in trans (R1E|H6A’) mutagenesis of R1 and H6 dramatically hinders Las1 RNA cleavage activity (Fig. 5B). Moreover, the detectable cleavage activity is predominantly the result of cleavage at the C2(−1) site. In accordance with our earlier work that shows the Las1 nuclease domain is important for supporting Grc3 kinase activity (23), we also observe a detectable RNA phosphorylation defect of the unprocessed RNA substrate for chimeric RNase PNK harboring Las1^R1E+H6A|WT’^. Our results reveal that Las1 cannot support catalysis with a single RHxhTH motif and that R1, R1’, H6 and H6’are all important for maintaining nuclease fidelity, likely through orienting the RNA substrate within the catalytic center. Therefore, these functional assays reveal residues within the composite active site that are critical for Las1 function and fidelity.

## Conclusion

In this study we report a comprehensive molecular characterization of the Las1 HEPN nuclease motif, which plays a critical role in pre-rRNA processing. HEPN nucleases participate in a wide spectrum of RNA processing pathways and the recent identification of many new HEPN nucleases (3) has led to a surge in atomic-resolution structures of different HEPN nuclease family members (Fig. 6) (4, 5, 26, 28–30, 35, 37–44). The overall architecture of HEPN nucleases varies dramatically from the butterfly-shaped RNase PNK to bilobed Cas13 nucleases and intertwined RNase L. These diverse assemblies are important contributors to the activation and specialized functions of individual HEPN nucleases. Yet, each of these nucleases contains a common homo- or hetero-dimeric HEPN core that is responsible for nuclease activity. The presence of this common core suggests that HEPN nucleases cleave RNA following a similar mechanism. Each HEPN core contains a composite nuclease active site at its center and is defined by the juxtaposition of two RϕxxxH motifs (Fig. 6). Despite numerous recent advances in HEPN nuclease biology, the molecular mechanism of RNA cleavage remains unresolved. By implementing an in-depth sequence analysis, yeast genetics and *in vitro* activity assays we determined that the first, second, fifth, and sixth residues from this motif are absolutely essential for supporting Las1 function. Due to the strong parallels amongst all HEPN nucleases, our work has broad implications for the HEPN nuclease field.

**Fig. 6.**
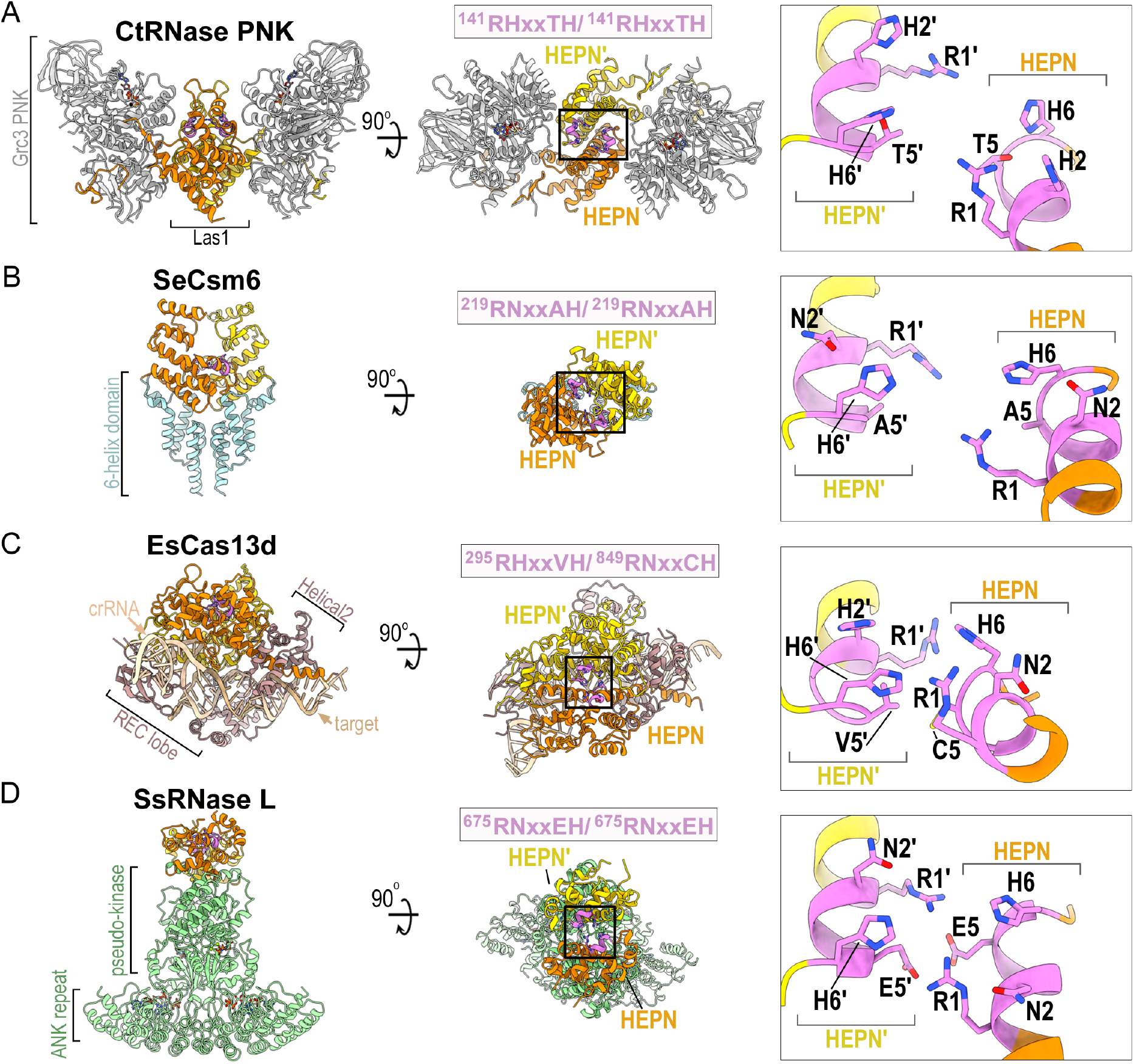
Structural comparison of HEPN nucleases. (*A*-*D*) Ribbon diagram of (*A*) *C. thermophilum* RNase PNK (PDB ID 6OF3), (*B*) *Staphylococcus epidermis* Csm6 (PDB ID 5yjc), (*C*) *Eubacterium siraeum* Cas13d (PDB ID 6e9f) and (*D*) *Sus scrofa* RNase L (PDB ID 4O1P). HEPN-HEPN’dimers are colored in orange and yellow along with their juxtaposed RϕxxxH motifs colored in purple. Enzyme-specific insertions of RNase PNK, Csm6, Cas13d and RNase L are colored in gray, light blue, brown and light green, respectively. Black boxes mark the HEPN nuclease active sites formed by well-conserved HEPN nuclease motifs (purple). The inset is a zoom of the juxtaposed RϕxxxH motifs forming the catalytic site. Conserved residues of the RϕxxxH motifs are shown and the second copy of the RϕxxxH motif is designated by prime.

### Significance of the Individual Residues from the HEPN Motif

While it’s well established that the canonical R1 and H6 residues from HEPN nucleases are important for catalysis (3, 7), this work highlights the significance of H2 and T5 from the Las1 HEPN motif. The invariant H6 residue is thought to be important for triggering 2’-OH nucleophilic attack, while R1 has been proposed to stabilize the transition state and/or the RNA substrate (3). Our work expands this working model by uncovering the functional significance of the intervening Las1 HEPN residues. Amino acid composition of the second residue from the HEPN motif is most often a polar residue. Indeed our complementation and *in vitro* cleavage assays revealed that Las1 can function with a conservative histidine to asparagine mutation at this position, but other substitutions are not tolerated. Similarly, HEPN nucleases Ire1, Csm6, and Cas13 encode an asparagine at its second position and previous work has shown that N2A mutants are functionally inactive (5, 27, 30). Together, this signifies a universal importance of the second residue from the RϕxxxH motif. Considering recent structural characterization of RNase PNK revealed that Las1 H2 is a functional molecular switch that undergoes conformational rearrangements within the HEPN active site, we compared the active sites from several recent HEPN nuclease structures. Intriguingly, within these structures the H2/N2 residue either points towards the center of the composite active site, as seen in the nuclease active state of RNase PNK, where it could coordinate RNA or it points away for the catalytic center, as seen in the nuclease inactive state of RNase PNK. This raises the question as to whether the second amino acid residue of the HEPN motif is a universal molecular switch regulating the nuclease activity of the HEPN superfamily. What then is the role of this residue? One possibility is that the H2 residue could participate in the general acid-base mechanism that has been proposed for HEPN family nucleases (3, 7, 36). However, this seems unlikely since we determined that H2N mutants of Las1 are functionally active, while H6N mutants are inactive. Furthermore, both asparagine and histidine can form hydrogen bonds with RNA, but only histidine is able to donate a proton because the pKa of asparagine is too high (45). Since the second position of the RϕxxxH motif is consistently a proton acceptor, we do not anticipate that H2 interacts with a phosphate group in the RNA backbone, but more likely interacts with a hydroxyl group on the pentose ring. Thus, we hypothesize that the second position residue of the HEPN motif is critical for proper positioning of the RNA substrate within the active site, while H6 participates in acid-base catalysis.

Our work also establishes the significance of T5 within the Las1 nuclease motif. Amongst HEPN nucleases there is a prevalence for a small amino acid at this position (2). This is observed in the recent structures of both Csm6 (A5/A5’) and Cas13d (V5/C5’), which harbor small amino acids at the fifth position (Fig. 6). Within composite HEPN nuclease active sites, the fifth position residue is located along the base of the active site. While the location of this residue suggests that it does not directly participate in catalysis, it likely plays a supportive role in maintaining the structural integrity of the active site. Here we show that Las1 is dependent upon having either a threonine or serine at this position. This dependence is supported by the sequence conversation amongst Las1 homologs and the presence of a hydrogen bond between T5 and H6 that was observed in the cryo-electron microscopy reconstruction of RNase PNK. Taken together, our work indicates that beyond the invariant R1 and H6 residues from the RϕxxxH HEPN nuclease motif, the intervening residues play important supporting roles in substrate engagement and active site architecture.

### It Takes Two (RϕxxxH Motifs) to Cut the RNA Right

A fundamental outstanding question is why are HEPN nucleases reliant on the juxtaposition of two RϕxxxH motifs to cleave the RNA backbone. A single RϕxxxH motif presumably contains the residues that are sufficient for cleavage, however all HEPN nucleases require dimerization to be active. There are several possibilities as to why dimerization could be important. For example, one RϕxxxH motif could be important for RNA cleavage while the other copy could participate in positioning the RNA. Another possibility is that the HEPN active site could be reliant on R1 from one motif and H6’from the other copy for catalysis. We suggest it is a combination of both since our work reveals that three of the four Las1 R1, R1’, H6 and H6’residues are needed for RNA hydrolysis. We also uncover through the creation of an RNase PNK chimera that Las1 strictly requires both RHxhTH motifs to properly cleave the ITS2 at the C2 site. Our characterization of a series of RNase PNK chimera variants revealed the surprising finding that several of the variants had altered cleavage specificity. The most striking alteration in specificity was observed with the WT|H6N’variant which cleaves the ITS2 predominantly at the C2(−1) position. In contrast, RNase L WT|H6N’chimeras, formed by mixing experiments, does not alter the cleavage specificity for its single stranded RNA substrate (28). An interesting possibility for the loss of specificity in RNase PNK and not RNase L could stem from the inherent difference in substrate selection employed by these two enzymes. RNase L displays broad specificity against many different RNA substrates and cleaves the RNA backbone between two residues following a uracil (UN^N) (28). This low specificity means the RNase L active site can accommodate numerous RNA substrates. On the other hand, RNase PNK has a single known RNA target, the ITS2 pre-rRNA. Minor modifications to the ITS2 pre-rRNA sequence and structure surrounding the C2 cleavage site abrogates cleavage *in vivo* and *in vitro* (23). Thus, while RNase PNK can cleave RNA with mutations in a single RHxhTH, it requires the cooperation of both RHxhTH motifs to ensure C2 cleavage specificity.

Defining the precise mechanism of HEPN-directed RNA cleavage will require a series of high resolution structures of HEPN domains engaged to their endogenous RNA substrates. However, to date the transient nature of RNA-associated HEPN nuclease states has proven challenging. Our work lays the foundation for understanding the contributions of the individual residues from the Las1 RHxhTH motif. Beyond advancing our understanding of pre-rRNA processing, the observation that Las1 can be re-wired to cleave RNA at different positions has far reaching implications in the rapidly developing field of RNA targeting applications. The Cas13 CRISPR effectors, which contain HEPN nuclease domains, are currently being adapted for a range of applications including RNA knockdown, RNA editing, and RNA detection/diagnostics (7–11, 46). Once activated by crRNA, the Cas13 HEPN domains orchestrate non-specific RNA cleavage. The ability to re-wire the Cas13 HEPN domains with altered specificity could open the door for new RNA-targeting applications.

## Materials and Methods

### Las1 Sequence Analysis

*Homo sapiens* Las1L (NP_112483.1) was used as a query to extract 101 Las1 orthologs for vertebrate genomes using Ensembl (www.ensembl.org, release 95; (47)) and *Saccharomyces cerevisiae* Las1 (NP_012989.3) was used as the query to extract 219 Las1 orthologs from Ensembl Fungi (release 42). Sequence alignments were performed with MAFFT 7 (https://www.ebi.ac.uk/Tools/msa/mafft/; (48)) using default settings. CLC Genomics version 8 was used for visualization of multiple sequence alignments and WebLogo (49) was used to generate the Las1 HEPN sequence logo.

### Generation of Las1 Yeast Strains

Generation of the *S. cerevisiae LAS1* tetracycline-titratable promoter (tetO_7_) strain was described previously (26). This strain was modified to include a 3xMyc-tag upstream of endogenous *GRC3* for detection purposes (yMP125). N-terminal 3xFlag-tagged ScLas1 (pMP 580) was amplified along with 300 nt of flanking endogenous DNA sequence and inserted into YCplac33 (32) using KpnI and SacI. Las1 HEPN variants were generated by overlap PCR, inserted into YCplac33 and verified by DNA sequencing (Genewiz). The yMP125 strain was transformed with plasmids encoding the N-terminal 3xFlag tagged Las1 wild type (pMP 580), Las1 HEPN variants (see Table S1), or empty YCplac33 vector. All yeast strains used in this study are listed in Table S2.

### Yeast Spotting Assays and Growth Curves

Yeast spotting assays and growth curves were performed as described previously (24) with minor modifications. Transformed *LAS1* yeast strains were pre-incubated in YPD that was supplemented with 40 μg/ml doxycycline for 24 hours at 22°C prior to performing assays. For proliferation assays, transformed *tet-LAS1* and *tet-LAS1/Myc-GRC3* strains (Table S2) were spotted on YPD plates in the absence and presence of doxycycline (40 μg/ml) and incubated at 30°C for 2-3 days. Temperature sensitivity was tested by carrying out additional proliferation assays at 16°C, 25°C, 34°C, and 37°C for 2-6 days. Growth curves were generated using transformed *tet-LAS1/Myc-GRC3* strains by measuring the absorbance at 595 nm of 100 μL yeast cultures inoculated at an OD_600_ of 0.05 and incubated at 30°C in YPD and YPD with doxycycline (40 μg/ml). OD_600_ measurements were recorded every 15 minutes over a 25 hour time period with an Infinite F200 Pro (Tecan) and i-control 1.11 software. The mean and standard deviation of each growth curve were calculated from three independent replicates.

### Western Blots

Western blotting was performed as described previously (24) with minor modifications. Transformed *tet-LAS1/Myc-GRC3* strains were grown in YPD in the presence of doxycycline (40 μg/ml) to mid-log phase. The whole cell lysate was prepared by lysing the cells with glass beads followed by trichloroacetic acid precipitation. Proteins were resolved by SDS-PAGE and analyzed by western blot using anti-Myc (Grc3; EMD Millipore), anti-Flag (Las1; Sigma), and anti-α-tubulin (Abcam).

### Recombinant Las1-Grc3 Complex Purification

*S. cerevisiae* RNase PNK variants were generated from the original Las1-Grc3 co-expression vector (pMP001) (23) by Q5 site-directed mutagenesis (NEB) and verified by DNA sequencing (Genewiz). ScLas1 HEPN|HEPN’fusion (Las1 residues 1-179 aa followed by a 2x Gly-Gly-Gly-Gly-Ser linker, NotI restriction site, and Las1 residues 1-185) was inserted into the expression plasmid ScLas1^LCT^-Grc3 (pMP072; (23)) using HindIII and SacI to generate ScLas1^HEPN|HEPN’+LCT^-Grc3 (pMP673). Chimeric ScLas1^HEPN|HEPN’+LCT^-Grc3 was generated by amplifying ScLas1 HEPN’(residues 1-185) harboring mutations R129E (R1E; pMP683), R129K (R1K; pMP684), H134A (H6A; pMP681) and H134N (H6N; pMP682) were amplified by PCR and inserted into pMP673 using NotI and SacI. Additional chimeric ScLas1^HEPN|HEPN’+LCT^-Grc3 constructs were generated by amplifying ScLas1 HEPN (residues 1-179) harboring mutations R129E+H134A (R1E+H6A; pMP697), H134A (H6A; pMP695) and R129E (R1E; pMP691) and inserted into pMP673 or pMP681 using HindIII and NotI. All Las1-Grc3 co-expression plasmids used and generated in this study were sequenced (GeneWiz) and are listed in Table S3.

Sc Las1-Grc3 variant proteins were expressed and purified as described previously (23) with minor modifications. Las1-Grc3 variants were overexpressed in *E. coli* LOBSTR (Kerafast) cells following cold shock and induction with 0.5 mM IPTG. Harvested cells were resuspended in lysis buffer (50 mM Tris pH 8.0, 500 mM NaCl, 5 mM MgCl_2_, 1%(v/v) Triton X-100, 10% (v/v) glycerol) and lysed by sonication. Clarified lysate was applied to a gravity flow column with His60 Ni Superflow Resin (Clontech) that was pre-equilibrated in lysis buffer. After washing with 200 mL of wash buffer (50 mM Tris pH 8.0, 500 mM NaCl, 5 mM MgCl_2_, 30 mM imidazole, 10% glycerol) the complex was eluted with imidazole (50 mM Tris pH 8.0, 500 mM NaCl, 5 mM MgCl_2_, 200 mM imidazole, 10% glycerol). Las1-Grc3 variants were further resolved by gel filtration (HiLoad 16/600 Superdex-200 Prep Grade; GE Healthcare) with storage buffer (20 mM Tris pH 8.0, 200 mM NaCl, 5 mM MgCl_2_, 5% glycerol).

### C2 pre-rRNA Cleavage and Phosphorylation Assays

C2 RNA cleavage assays were performed as described previously (23, 24) with minor changes. Full-length ScLas1-Grc3 and chimeric ScLas1^HEPN|HEPN’+LCT^-Grc3 variants (0-3.2 μM) were incubated with 1 mM EDTA and 50 or 100 nM 3’-fluorescein labeled C2 RNA substrate (5’-GUCGUUUUAGGUUUUACCAACUGCGGC/36-FAM/-3’) in the absence and presence of 2 mM nucleotide (ATP or ADPnP). Cleavage reactions were incubated for 1 hour at 37°C and then quenched with urea-loading dye (20 mM Tris pH 8.0, 8 M urea, 0.05% bromophenol blue, 1 mM EDTA). Samples were resolved on 15% polyacrylamide (8 M urea) gels in 1x tris-borate-EDTA buffer and visualized using a Typhoon FLA 9500. Representative gels are shown from three independent replicates.

### Mass Spectrometric Sample Preparation and Analysis of Oligonucleotides

10 μL of RNA sample (5 μM) was injected onto the column for liquid chromatography mass spectrometry (LC-MS) analysis. Data were acquired on a Q Exactive Plus mass spectrometer (QE-MS, ThermoFisher Scientific) interfaced with a Vanquish (ThermoFisher Scientific) UHPLC system. Reverse-phase chromatography was performed using a CORTECS C18 column (100 x 2.1 mm i.d., 1.6 μm particle size; Waters Corporation) and a CORTECS C18 VanGuard precolumn (5 x 2.1 mm i.d., 1.6 μm particle size; Waters Corporation) with solvent A being 5 mM ammonium formate in water (pH 6.5) and solvent B being methanol and a flow rate of 150 μL per minute. The LC gradient included a ramp from 0% to 42% B from 0 to 6 minutes followed by a ramp from 42% to 95% B over one minute and then a 3-minute hold at 95% B. The run was completed with a ramp of 95% to 0% B for 0.5 minutes followed by a 4.5 minute recondition at 0% B. The QE-MS was equipped with a HESI source used in the negative ion mode and performing only MS1 scans. Mass calibration was performed before data acquisition using the Pierce ESI Negative Ion Calibration Mixture (Pierce). Data were processed and deconvoluted using the Intact Protein Analysis function of BioPharma Finder (ThermoFisher Scientific). Mass predictions of the oligonucleotides were performed using the web version of Mongo Oligo Mass Calculator v2.08 maintained by the RNA Institute.

## Acknowledgements

We thank Drs. Lars Pedersen and Chen Qiu, as well as all the members of the Stanley Lab for their critical reading of this manuscript. We are grateful to Dr. Thomas Randall from the NIEHS Integrative Bioinformatics Support Group for his help with Las1 sequence alignments. This work was supported by the US National Institute of Health Intramural Research Program; US National Institute of Environmental Health Sciences (NIEHS; ZIA ES103247 to R.E.S) and the Canadian Institutes of Health Research (CIHR; 146626 to M.C.P).

---

**Fig. S1.**
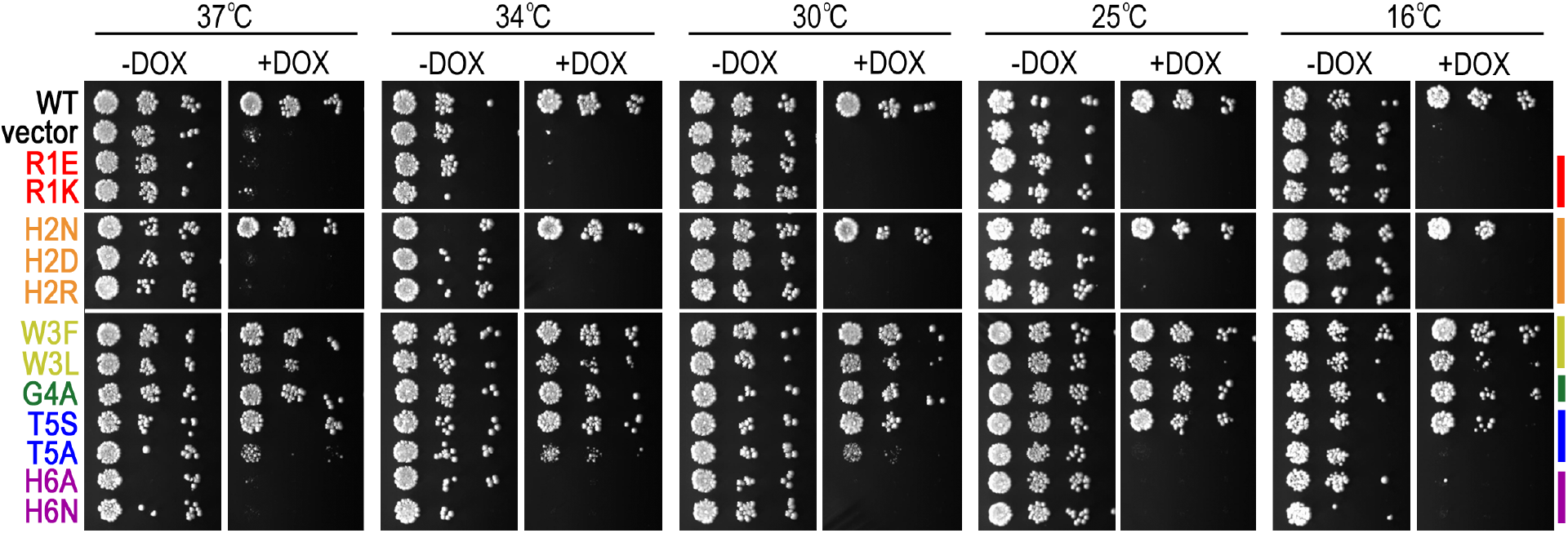
*In vivo* assay for complementation of *S. cerevisiae LAS1* over a temperature range. The *S. cerevisiae tet-LAS1/3xMyc-GRC3* strain was transformed with plasmids encoding Flag-Las1 RHxhTH variants (see Table S2). Serial dilutions were spotted on YPD in the absence (-DOX) and presence (+DOX) of 40 μg/ml doxycycline and incubated at 37°C, 34°C, 30°C, 25°C and 16°C for 2-6 days.

**Fig. S2.**
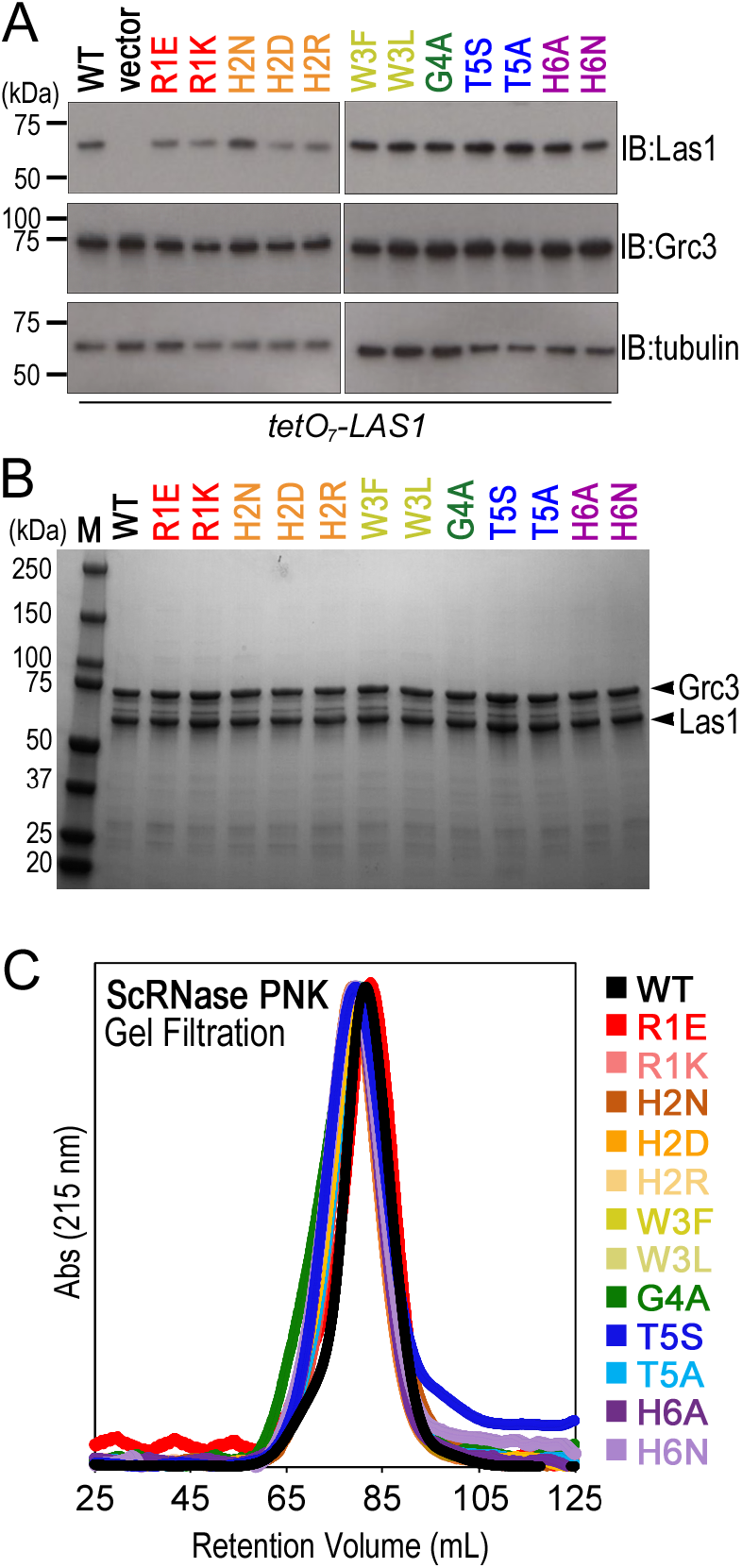
Las1 RHxhTH variants retain its association to Grc3. (*A*) Transformed *S. cerevisiae tet-LAS1/3xMyc-GRC3* strains were grown to mid-log phase at 30°C in the presence of 40 μg/ml doxycycline. Cells were lysed and equal amounts of whole cell lysate were separated by SDS-PAGE and analyzed by western blotting with anti-Flag (Las1), anti-Myc (Grc3) and anti-tubulin (loading control). IB is an abbreviation for immunoblotting. (*B*) SDS-PAGE analysis of purified recombinant ScRNase PNK complex comprised of full length Grc3 and Las1 RHxhTH variants. (*C*) Gel filtration of ScRNase PNK variants containing Las1 RHxhTH missense mutants. Wild type RNase PNK (black) was previously shown to form a hetero-tetramer (23) and has the same retention volume as the recombinant ScRNase PNK variants. This suggests that the ScRNase PNK variants maintain higher-order assembly.

**Fig. S3.**
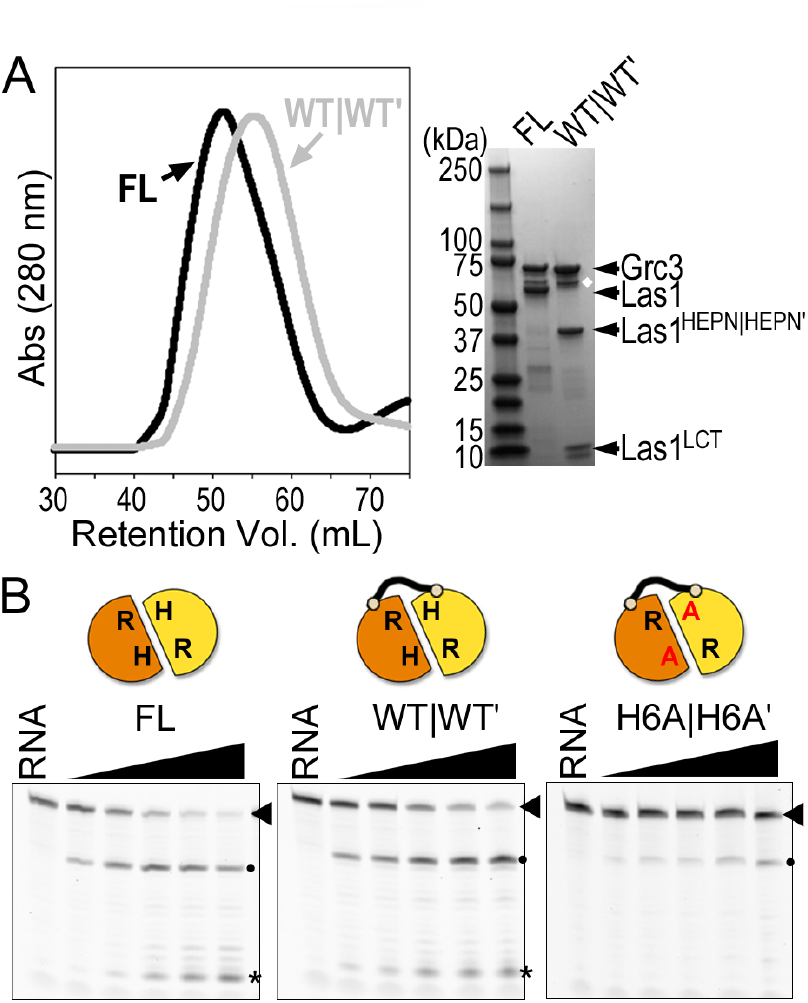
Functional *S. cerevisiae* RNase PNK chimera for *in vitro* assays. (*A*) Gel filtration and SDS-PAGE analysis of full length ScRNase PNK (FL; black), composed of Las1 and Grc3, and chimeric ScRNase PNK (WT|WT’; gray), comprised of Las1 HEPN|HEPN, Las1 LCT and Grc3. White diamond marks a Grc3 degradation product. Chimeric ScRNase PNK has a similar retention volume to the full length complex suggesting the chimera maintains its higher-order assembly despite lacking its CC domain. (*B*) Denaturing urea gels of reactions containing chimeric Sc RNase PNK variants (0-3.2 μM) encoding missense mutations to the Las1 RHxhTH motif. Protein variants were incubated with C2 RNA substrate (50 nM) for 1 hour at 37°C. RNA defines reactions set in the absence of protein and asterisks marks the accumulation of a non-specific band that is dependent on the presence of active RNase PNK complex and a 3’-fluorescent label. Dots mark RNA cleavage products where black dots are the result of cleavage at the C2 site and red dots are off-target cleavage events.

**Fig. S4.**
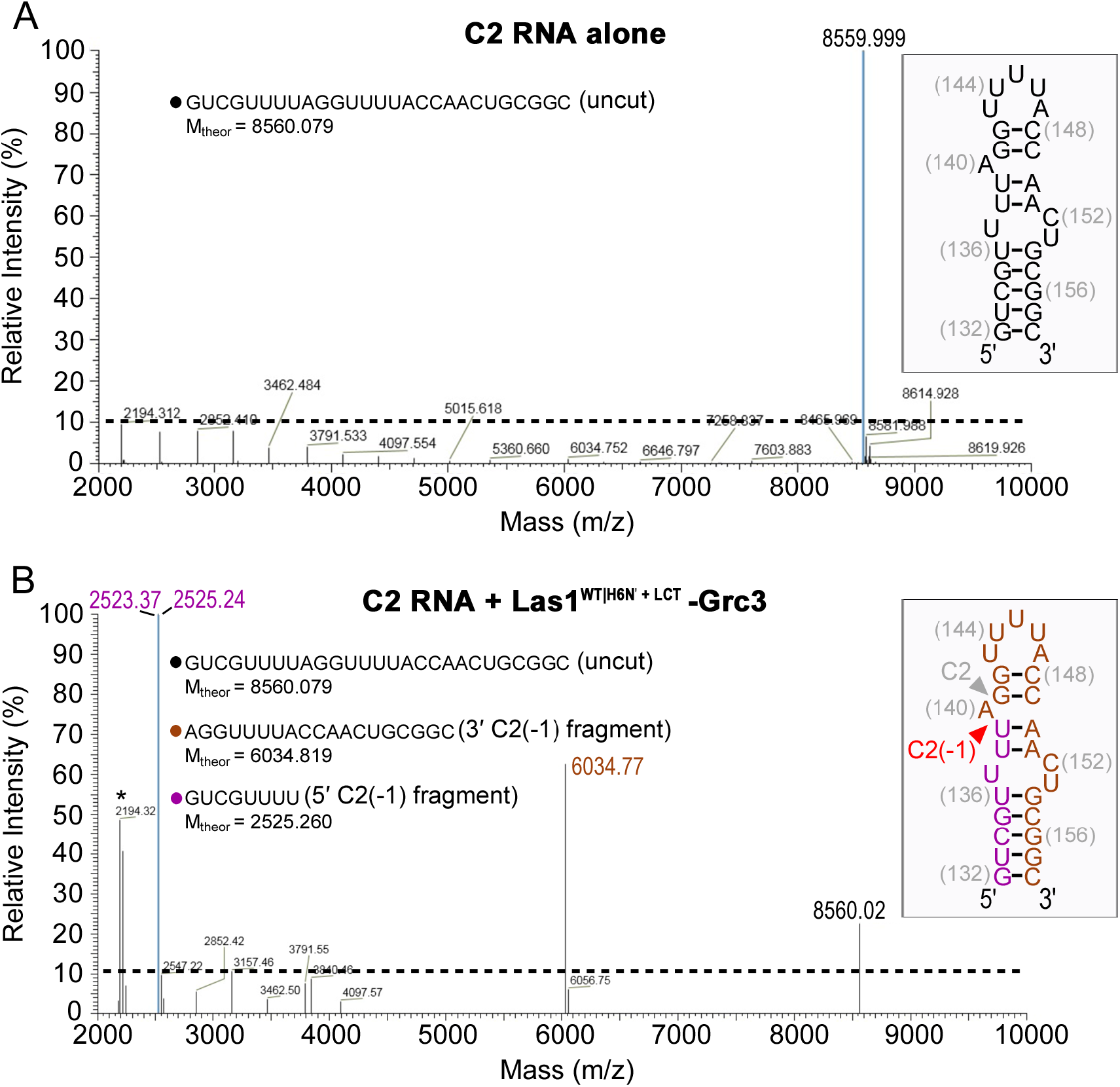
Identification of off-target RNA products using mass spectrometry. (*A-B*) Spectra by LC-ESI-MS of C2 RNA substrate incubated in the (*A*) absence and (*B*) presence of chimeric RNase PNK containing Las1^WT|H6N’^. RNA (10 μM) was incubated in the absence and presence of protein (13 μM) for 30 minutes at 37°C. Observed ions of *m/z* 8560 is indicative of the uncut C2 RNA substrate. Whereas ions of *m/z* 2525.3 and 6034.8 correspond to 5’- and 3’-RNA fragments, respectively, produced following cleavage at an off-target site (C2(−1); red arrowhead) between U139 and A140 of the *S. cerevisiae* ITS2. The 8-nucleotide 5’-fragment (magenta) harbors a 2’,3’-cyclic phosphate and the 19-nucleotide 3’-fragment (brown) contains a 5’-hydroxyl end confirming the ScLas1^WT|H6N’^ variant is responsible for the off-target RNA hydrolysis.

**Table S1.** Yeast plasmids used in this study

| Plasmid | Variant | *LAS1* | Vector | Source |
| --- | --- | --- | --- | --- |
| pMP580 | ScLas1 | WT; residues 1-502,<br>3x N-terminal FLAG Tag | YCplac33 | (26) |
| pMP600 | ScLas1<sup>R1E</sup> | R129E; residues 1-502,<br>3x N-terminal FLAG Tag | YCplac33 | This study |
| pMP648 | ScLas1<sup>R1K</sup> | R129K; residues 1-502,<br>3x N-terminal FLAG Tag | YCplac33 | This study |
| pMP632 | ScLas1<sup>H2N</sup> | H130N; residues 1-502,<br>3x N-terminal FLAG Tag | YCplac33 | This study |
| pMP631 | ScLas1<sup>H2D</sup> | H130D; residues 1-502,<br>3x N-terminal FLAG Tag | YCplac33 | This study |
| pMP654 | ScLas1<sup>H2R</sup> | H130R; residues 1-502,<br>3x N-terminal FLAG Tag | YCplac33 | This study |
| pMP601 | ScLas1<sup>W3F</sup> | W131F; residues 1-502,<br>3x N-terminal FLAG Tag | YCplac33 | This study |
| pMP602 | ScLas1<sup>W3L</sup> | W131L; residues 1-502,<br>3x N-terminal FLAG Tag | YCplac33 | This study |
| pMP603 | ScLas1<sup>G4A</sup> | G132A; residues 1-502,<br>3x N-terminal FLAG Tag | YCplac33 | This study |
| pMP590 | ScLas1<sup>T5S</sup> | T133S; residues 1-502,<br>3x N-terminal FLAG Tag | YCplac33 | This study |
| pMP656 | ScLas1<sup>T5A</sup> | T133A; residues 1-502,<br>3x N-terminal FLAG Tag | YCplac33 | This study |
| pMP591 | ScLas1<sup>H6A</sup> | H134A; residues 1-502,<br>3x N-terminal FLAG Tag | YCplac33 | This study |
| pMP649 | ScLas1<sup>H6N</sup> | H134N; residues 1-502,<br>3x N-terminal FLAG Tag | YCplac33 | This study |

**Table S2.** Yeast strains used and constructed in this study

| Strain | Genotype | Source |
| --- | --- | --- |
| *tet-LAS1* (CML476) | *MATα*; *ura3-52*; *leu2Δ1*; *his3Δ200*; *GAL2*; *CMVp(tetR'-SSN6)::LEU2*; *trp1::tTa* | EUROSCARF |
| *tet-LAS1* + *pMP580* | *MATα*; *ura3-52*; *leu2Δ1*; *his3Δ200*; *GAL2*; *CMVp(tetR'-SSN6)::LEU2*; *trp1::tTa*; *pMP 580 (3xFlag-LAS1; WT)* | This Study |
| *tet-LAS1/Myc-Grc3* (yMP125) | *MATα*; *ura3-52*; *leu2Δ1*; *his3Δ200*; *GAL2*; *CMVp(tetR'-SSN6)::LEU2*; *trp1::tTa*; *kanMX:tetO<sub>7</sub>:LAS1*; *trp1:3xMyc:GRC3* | (26) |
| *tet-LAS1/Myc-Grc3* + *pMP580* | *MATα*; *ura3-52*; *leu2Δ1*; *his3Δ200*; *GAL2*; *CMVp(tetR'-SSN6)::LEU2*; *trp1::tTa*; *kanMX:tetO<sub>7</sub>:LAS1*; *trp1:3xMyc:GRC3*; *pMP 580 (3xFlag-LAS1; WT)* | (26) |
| *tet-LAS1/Myc-Grc3* + *pMP600* | *MATα*; *ura3-52*; *leu2Δ1*; *his3Δ200*; *GAL2*; *CMVp(tetR'-SSN6)::LEU2*; *trp1::tTa*; *kanMX:tetO<sub>7</sub>:LAS1*; *trp1:3xMyc:GRC3*; *pMP 600 (3xFlag-LAS1; R129E)* | This Study |
| *tet-LAS1/Myc-Grc3* + *pMP648* | *MATα*; *ura3-52*; *leu2Δ1*; *his3Δ200*; *GAL2*; *CMVp(tetR'-SSN6)::LEU2*; *trp1::tTa*; *kanMX:tetO<sub>7</sub>:LAS1*; *trp1:3xMyc:GRC3*; *pMP 648 (3xFlag-LAS1; R129K)* | This Study |
| *tet-LAS1/Myc-Grc3* + *pMP632* | *MATα*; *ura3-52*; *leu2Δ1*; *his3Δ200*; *GAL2*; *CMVp(tetR'-SSN6)::LEU2*; *trp1::tTa*; *kanMX:tetO<sub>7</sub>:LAS1*; *trp1:3xMyc:GRC3*; *pMP 632 (3xFlag-LAS1; H130N)* | This Study |
| *tet-LAS1/Myc-Grc3* + *pMP631* | *MATα*; *ura3-52*; *leu2Δ1*; *his3Δ200*; *GAL2*; *CMVp(tetR'-SSN6)::LEU2*; *trp1::tTa*; *kanMX:tetO<sub>7</sub>:LAS1*; *trp1:3xMyc:GRC3*; *pMP 631 (3xFlag-LAS1; H130D)* | This Study |
| *tet-LAS1/Myc-Grc3* + *pMP654* | *MATα*; *ura3-52*; *leu2Δ1*; *his3Δ200*; *GAL2*; *CMVp(tetR'-SSN6)::LEU2*; *trp1::tTa*; *kanMX:tetO<sub>7</sub>:LAS1*; *trp1:3xMyc:GRC3*; *pMP 654 (3xFlag-LAS1; H130R)* | This Study |
| *tet-LAS1/Myc-Grc3* + *pMP601* | *MATα*; *ura3-52*; *leu2Δ1*; *his3Δ200*; *GAL2*; *CMVp(tetR'-SSN6)::LEU2*; *trp1::tTa*; *kanMX:tetO<sub>7</sub>:LAS1*; *trp1:3xMyc:GRC3*; *pMP 601 (3xFlag-LAS1; W131F)* | This Study |
| *tet-LAS1/Myc-Grc3* + *pMP602* | *MATα*; *ura3-52*; *leu2Δ1*; *his3Δ200*; *GAL2*; *CMVp(tetR'-SSN6)::LEU2*; *trp1::tTa*; *kanMX:tetO<sub>7</sub>:LAS1*; *trp1:3xMyc:GRC3*; *pMP 602 (3xFlag-LAS1; W131L)* | This Study |
| *tet-LAS1/Myc-Grc3* + *pMP603* | *MATα*; *ura3-52*; *leu2Δ1*; *his3Δ200*; *GAL2*; *CMVp(tetR'-SSN6)::LEU2*; *trp1::tTa*; *kanMX:tetO<sub>7</sub>:LAS1*; *trp1:3xMyc:GRC3*; *pMP 603 (3xFlag-LAS1; G132A)* | This Study |
| *tet-LAS1/Myc-Grc3* + *pMP590* | *MATα*; *ura3-52*; *leu2Δ1*; *his3Δ200*; *GAL2*; *CMVp(tetR'-SSN6)::LEU2*; *trp1::tTa*; *kanMX:tetO<sub>7</sub>:LAS1*; *trp1:3xMyc:GRC3*; *pMP 590 (3xFlag-LAS1; T133S)* | This Study |
| *tet-LAS1/Myc-Grc3* + *pMP656* | *MATα*; *ura3-52*; *leu2Δ1*; *his3Δ200*; *GAL2*; *CMVp(tetR'-SSN6)::LEU2*; *trp1::tTa*; *kanMX:tetO<sub>7</sub>:LAS1*; *trp1:3xMyc:GRC3*; *pMP 656 (3xFlag-LAS1; T133A)* | This Study |
| *tet-LAS1/Myc-Grc3* + *pMP591* | *MATα*; *ura3-52*; *leu2Δ1*; *his3Δ200*; *GAL2*; *CMVp(tetR'-SSN6)::LEU2*; *trp1::tTa*; *kanMX:tetO<sub>7</sub>:LAS1*; *trp1:3xMyc:GRC3*; *pMP 591 (3xFlag-LAS1; H134A)* | This Study |
| *tet-LAS1/Myc-Grc3* + *pMP649* | *MATα*; *ura3-52*; *leu2Δ1*; *his3Δ200*; *GAL2*; *CMVp(tetR'-SSN6)::LEU2*; *trp1::tTa*; *kanMX:tetO<sub>7</sub>:LAS1*; *trp1:3xMyc:GRC3*; *pMP 649 (3xFlag-LAS1; H134N)* | This Study |

**Table S3.** *E. coli* expression plasmids used and constructed in this study

| Plasmid | Variant | *GRC3* | *LAS1* | Vector | Source |
| --- | --- | --- | --- | --- | --- |
| pMP001 | ScLas1-Grc3 | 1-632 aa | 1-502 aa | pST39 | (23) |
| pMP538 | ScLas1<sup>R1E</sup>-Grc3 | 1-632 aa | 1-502 aa; R129E | pST39 | This Study |
| pMP593 | ScLas1<sup>R1K</sup>-Grc3 | 1-632 aa | 1-502 aa; R129K | pST39 | This Study |
| pMP531 | ScLas1<sup>H2N</sup>-Grc3 | 1-632 aa | 1-502 aa; H130N | pST39 | This Study |
| pMP530 | ScLas1<sup>H2R</sup>-Grc3 | 1-632 aa | 1-502 aa; H130R | pST39 | This Study |
| pMP594 | ScLas1<sup>H2D</sup>-Grc3 | 1-632 aa | 1-502 aa; H130D | pST39 | This Study |
| pMP629 | ScLas1<sup>W3F</sup>-Grc3 | 1-632 aa | 1-502 aa; W131F | pST39 | This Study |
| pMP628 | ScLas1<sup>W3L</sup>-Grc3 | 1-632 aa | 1-502 aa; W131L | pST39 | This Study |
| pMP541 | ScLas1<sup>G4A</sup>-Grc3 | 1-632 aa | 1-502 aa; G132A | pST39 | This Study |
| pMP517 | ScLas1<sup>T5S</sup>-Grc3 | 1-632 aa | 1-502 aa; T133S | pST39 | This Study |
| pMP595 | ScLas1<sup>T5A</sup>-Grc3 | 1-632 aa | 1-502 aa; T133A | pST39 | This Study |
| pMP539 | ScLas1<sup>H6A</sup>-Grc3 | 1-632 aa | 1-502 aa; H134A | pST39 | This Study |
| pMP592 | ScLas1<sup>H6N</sup>-Grc3 | 1-632 aa | 1-502 aa; H134N | pST39 | This Study |
| pMP673 | ScLas1<sup>WT\|WT'+LCT</sup>-Grc3 | 1-632 aa | HEPN-HEPN': 1-179 aa, 2x GGGGS linker, 1-185 aa + LCT: 469-502 aa | pST39 | This Study |
| pMP681 | ScLas1<sup>WT\|H6A'+LCT</sup>-Grc3 | 1-632 aa | HEPN-HEPN': 1-179 aa, 2x GGGGS linker, 1-185 aa (H134A) + LCT: 469-502 aa | pST39 | This Study |
| pMP682 | ScLas1<sup>WT\|H6N'+LCT</sup>-Grc3 | 1-632 aa | HEPN-HEPN': 1-179 aa, 2x GGGGS linker, 1-185 aa (H134N) + LCT: 469-502 aa | pST39 | This Study |
| pMP683 | ScLas1<sup>WT\|R1E'+LCT</sup>-Grc3 | 1-632 aa | HEPN-HEPN': 1-179 aa, 2x GGGGS linker, 1-185 aa (R129E) + LCT: 469-502 aa | pST39 | This Study |
| pMP684 | ScLas1<sup>WT\|R1K'+LCT</sup>-Grc3 | 1-632 aa | HEPN-HEPN': 1-179 aa, 2x GGGGS linker, 1-185 aa (R129K) + LCT: 469-502 aa | pST39 | This Study |
| pMP691 | ScLas1<sup>R1E\|H6A'+LCT</sup>-Grc3 | 1-632 aa | HEPN-HEPN': 1-179 aa (R129E), 2x GGGGS linker, 1-185 aa (H134A) + LCT: 469-502 aa | pST39 | This Study |
| pMP697 | ScLas1<sup>R1E,H6A\|WT'+LCT</sup>-Grc3 | 1-632 aa | HEPN-HEPN': 1-179 aa (R129E, H134A), 2x GGGGS linker, 1-185 aa + LCT: 469-502 aa | pST39 | This Study |
| pMP695 | ScLas1<sup>H6A\|H64A'+LCT</sup>-Grc3 | 1-632 aa | HEPN-HEPN': 1-179 aa (H134A), 2x GGGGS linker, 1- | pST39 | This Study |
|  |  |  | 185 aa (H134A) + LCT: 469-502<br>aa |  |  |

